# Genome assembly of three Amazonian *Morpho* butterfly species reveals Z-chromosome rearrangements between closely-related species living in sympatry

**DOI:** 10.1101/2022.10.26.513852

**Authors:** Héloïse Bastide, Manuela López-Villavicencio, David Ogereau, Joanna Lledo, Anne-Marie Dutrillaux, Vincent Debat, Violaine Llaurens

**Affiliations:** IDEEV, Bât. 680,12, route 128, 91190 Gif Sur Yvette, France; Institut de Systématique, Evolution et Biodiversité (UMR 7205 CNRS/MNHN/SU/EPHE/UA), Muséum National d’Histoire Naturelle - CP50, 45 rue Buffon, 75005 PARIS, France; GeT-PlaGe, Bât G2, INRAe, 24 chemin de borde rouge - Auzeville, CS 52627, 31326 CASTANET-TOLOSAN Cedex, France

## Abstract

The genomic processes enabling speciation and the coexistence of species in sympatry are still largely unknown. Here we describe the whole genome sequencing and assembly of three closely-related species from the butterfly genus *Morpho*: *Morpho achilles* (Linnaeus, 1758), *M. helenor* (Cramer, 1776) and *M. deidamia* (Hübner, 1819). These large blue butterflies are emblematic species of the Amazonian rainforest. They live in sympatry in a wide range of their geographical distribution and display parallel diversification of dorsal wing colour pattern, suggesting local mimicry. By sequencing, assembling and annotating their genomes, we aim at uncovering pre-zygotic barriers preventing gene flow between these sympatric species. We found a genome size of 480 Mb for the three species and a chromosomal number ranging from 2n = 54 for *M. deidamia* to 2n = 56 for *M. achilles* and *M. helenor*. We also detected inversions on the sex chromosome Z that were differentially fixed between species, suggesting that chromosomal rearrangements may contribute to their reproductive isolation. The annotation of their genomes allowed us to recover in each species at least 12,000 protein-coding genes and to discover duplications of genes potentially involved in pre-zygotic isolation like genes controlling colour discrimination (*L-opsin*). Altogether, the assembly and the annotation of these three new reference genomes open new research avenues into the genomic architecture of speciation and reinforcement in sympatry, establishing *Morpho* butterflies as a new eco-evolutionary model.

## Introduction

Chromosomal rearrangements are likely to play a major role in both adaptation and speciation processes ([1, 2]). Inversions, for instance, can favour the emergence of adaptive syndromes by locking together co-adapted allelic variations [3]. Chromosomal rearrangements have also been suggested to contribute to reproductive isolation between species by promoting divergent adaptation or by bringing together genetic incompatibilities [4]. Nevertheless, the role of structural variants in these evolutionary processes is still largely unknown. Recently-developed sequencing and assembly methods now provide access to complete genomes, therefore opening the investigation of structural variation within and among species (see [5] for a review).

Here, we focus on emblematic species of the Amazonian rainforest, the blue *Morpho*. We describe the whole genome sequences of three closely-related *Morpho* species living in sympatry for a large range of their geographical distribution (Fig. 1): *M. helenor, M. achilles* and M. *deidamia* [6], thereby developing relevant resources to study the evolution of barriers to gene flow in sympatry. In Lepidoptera, specialization towards host-plant has been shown to be a major factor affecting species diversification [7]. Such ecological specialization may favour speciation and co-existence in sympatry, and may stem from the evolution of gustatory receptors enabling plant recognition by females [8].

**Figure 1.**
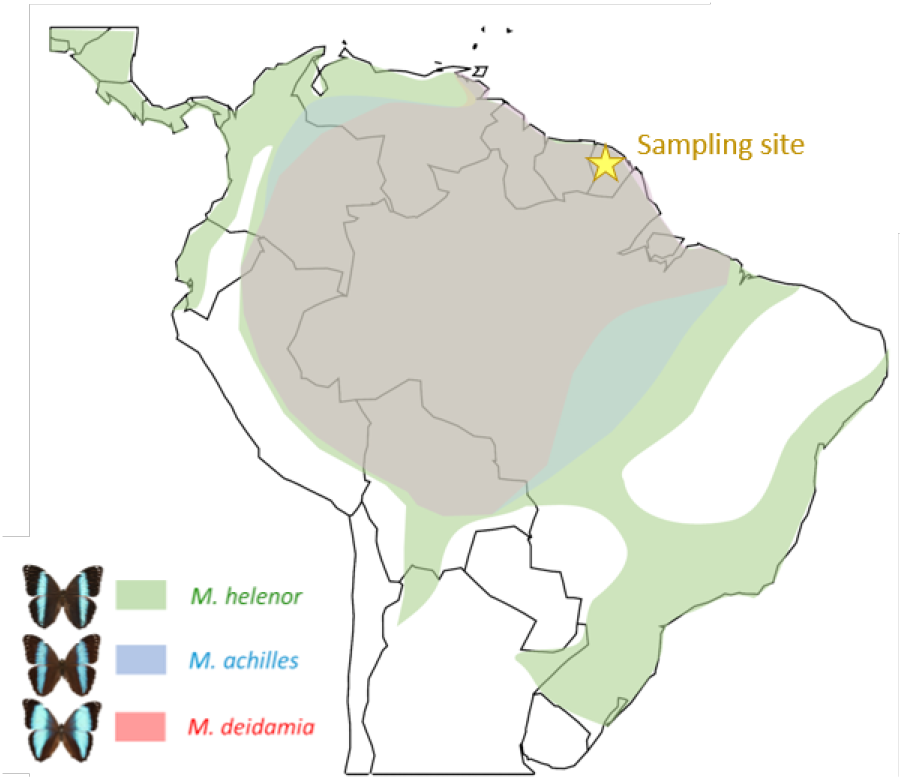
Geographical distribution of the three neotropical species *M. helenor* (green areas), M. *achilles* (blue areas) and M. *deidamia* (red areas). *M. helenor* has the widest distribution, from central America to Southern Brazil, while *M. achilles* and *M. deidamia* are restricted to the Amazonian basin. The three species are in sympatry throughout the Amazonian rainforest, including French Guiana (marked with the yellow star) where the samples studied here were collected.

The evolution of visual [9] and olfactory signals [10] between species may also limit gene flow between sympatric species of Lepidoptera. In the three *Morpho* species studied here, both males and females display conspicuous iridescent blue colour patterns on the dorsal side of their wings, combined with cryptic brownish colour on the ventral side [11]. Such a combination of dorso-ventral pattern, associated with a fast and erratic flight, is thought to contribute to the high escape abilities from predators of these butterflies, promoting colour pattern convergence between sympatric species (*i.e*. escape mimicry, [12]). Parallel geographic variation of dorsal wing colour pattern has indeed been detected in the three *Morpho* species studied here, suggesting local convergence promoted by predators behaviour [13]. Given the key role of colour pattern in both sexual selection and species recognition in diurnal butterflies, such a resemblance is thought to enhance reproductive interference between sympatric species [14]. Behavioural experiments carried out in the wild revealed that males from the three mimetic *Morpho* species are indeed attracted by both intra and interspecific wing patterns [15]. Despite this heterospecific attraction of males at long distances, RAD-sequencing markers revealed a highly limited gene flow between these three sympatric species [15]. This might be due to the differences in the timing of daily activities observed between these sympatric species limiting heterospecific encountering [15]. This divergence in daily phenology may contribute to the initiation of speciation or to the reinforcement of pre-zygotic barriers to heterospecific matings.

Genetic incompatibilities may also contribute to speciation and reinforcement processes by generating post-zygotic barriers. For instance, variation in chromosome numbers has been shown to correlate with speciation rate in Lepidoptera [16]. Similarly, chromosomal inversions may fuel the speciation process: by capturing genetic variations, inversions may lead to increased genetic divergence between species. Such divergence may lead to maladaption in hybrids and further limit gene flow between species living in sympatry.

By relying on both karyotype data and PacBio-Hifi sequencing, we generated *de novo* genome assemblies for three sympatric species of *Morpho* butterflies. The divergence between the two sister species *M. helenor* and *M. achilles* was estimated to occur about 3.91 My ago, while the divergence between these two sister species and *M. deidamia* was estimated to circa 16.68 My ago [17], enabling to compare the genome divergence in sympatry at different time scales. We then investigated the structural variants and variation in genes potentially contributing to pre-zygotic isolation among these species. We aim to shed light on the genomic processes involved in sympatric speciation and reinforcement as well as detecting chromosomal rearrangements. We also provide their mitochondrial genomes, study their transposable element (TE) contents and annotate the genomes. These genomic resources will open new research avenues into the understanding of adaptive processes, such as convergence evolution of colour pattern or divergence in visual systems, as well as speciation and co-existence of sister-species in sympatry, establishing *Morpho* butterflies as a new eco-evolutionary model

## Materials and Methods

### Butterfly sampling

Males from the species *M. helenor* (*n* = 1), *M. achilles* (*n* = 4) and *M. deidamia* (*n* = 2) were caught with a handnet at the Patawa waterfall, located in the Kaw mountain area of French Guiana (GPS location: 4.54322; −52.15832) to perform DNA extractions. In these species, males typically patrol in river beds and are easy to catch, while females are more rarely encountered. We therefore focused on males only. Because in butterflies sex is controlled by a ZW sex chromosome system (females being the heterogametic sex), we were thus able to access the Z sex chromosome but not the W chromosome.

### Karyotype study

Cytogenetic techniques were applied to wild caught males (*M. helenor* (*n* = 3), *M. achilles* (*n* = 4) and *M. deidamia* (*n* = 2)) that were collected at the abovementioned location in 2019. Their testicles were dissected and processed shortly after capture following the protocol described in [18]. The obtained cell suspension was conserved in fixative at about 4°C. The cell spreading and staining were then performed as described in [18]. The chromosome staining relied on the Giemsa method.

### DNA extractions and genome sequencing

Live butterflies (*M. helenor* (*n* = 1), *M. achilles* (*n* = 4) and *M. deidamia* (*n* = 2)) captured in 2021 at the same site in French Guiana were killed in the lab and their body immediately placed in liquid nitrogen. The DNA extraction was carried out the following day using the Qiagen Genomic-tip 100/G kit and following supplier instructions. The extracted DNA of a single male from each species was used (see Supplementary Fig. 1 for pictures of the wings of the sequenced specimens). Library preparation and sequencing were performed at GeT-PlaGe core facility (INRAe Toulouse) according to the manufacturer’s instructions “Procedure and Checklist Preparing HiFi SMRTbell Libraries using SMRTbell Express Template Prep Kit 2.0”. At each step, DNA was quantified using the Qubit dsDNA HS Assay Kit (Life Technologies). DNA purity was tested using the nanodrop (Thermofisher) and size distribution and degradation assessed using the Femto pulse Genomic DNA 165 kb Kit (Agilent). Purification steps were performed using AMPure PB beads (Pacific Biosciences). 15μg of DNA was purified then sheared at 15kb (speed 31 and 32) with the Megaruptor3 system (Diagenode). Using SMRTbell Express Template prep kit 2.0, a Single strand overhangs removal, a DNA and END damage repair step were performed on 10μg of sample. Blunt hairpin adapters were then ligated to the library, which was treated with an exonuclease cocktail to digest unligated DNA fragments. A size selection step using a 10kb cutoff was performed on the BluePippin Size Selection system (Sage Science) with the “0.75 percent DF Marker S1 6-10 kb vs3 Improved Recovery” protocol. Using Binding kit 2.2 and sequencing kit 2.0, the primer V5 annealed and polymerase 2.2 bounded library was sequenced by diffusion loading onto 1 SMRTcells per sample on SequelII instrument at 80 pM with a 2 hours pre-extension and a 30 hours movie.

### K-mer analysis, genome size and heterozygosity estimation

We used Jellyfish (v.2.3.0) [19] to perform a k-mer analysis on each PacBio dataset with a *k*-mer size of 21. For each dataset *k*-mers were counted and aggregated (jellyfish count option) and histograms were generated using the “-histo” command. The resulting histograms allowed the estimation of genome length and heterozygosity with GenomeScope version 2.0 [20] using the web application.

### Nuclear and mitochondrial genome assembly

For the assembly of the nuclear genomes, we compared three long-read assembly tools: IPA-Improved Phased Assembler (v1.0.3-0) (https://github.com/PacificBiosciences/-pbipa, Flye (v2.9) [21] and Hifiasm (v0.16.1 with the option -l3 to purge all types of haplotigs in the most aggressive way) [22]. For each assembler, we estimated basic assembly statistics such as scaffold count, contig count and N50 using the “stats.sh” program from the BBMap v38.93 package [23]. The completeness of each assembly was assessed using BUSCO v5.2.2 and MetaEuk for gene prediction against the *lepidoptera_odb10* database [24]. We retained the Hifiasm assembly because it had the highest BUSCO score, the highest contiguity (N50) and longest contig. Despite the high level of purging performed by Hifiasm, the species (*M. helenor* and *M. achilles* respectively) retained a high level of duplicates in the BUSCO score. To remove false haplotypic duplications in these two species, we used Purge_dups v1.2.5 setting the cutoffs manually (with calcuts -l 5 -m 33 -u 135 for *M. helenor* and calcuts -l 10 -m 45 -u 145 for *M. achilles*) [25]. The completeness of the purged genomes was then reassessed using BUSCO.

The mitochondrial genome of each species was assembled and circularized using Rebaler (https://github.com/rrwick/Rebaler) directly from the PacBio Hifi reads and using the mitochondrial genome of the closely related species *Pararge aegeria* as a reference.

### Annotation of repetitive regions

The annotation of repetitive regions in the three species was performed following two main steps. First, we used RepeatModeler v2.0.2a [26] with the option -s (slow search) and -a (to get a .align output file) to create *de novo* libraries of repetitive elements for each species. The library was then used to hardmask the corresponding genome assembly using RepeatMasker 4.1.2.p1 [26] . A summary of the repeated elements was generated with the script ‘buildSummary.pl’ included in RepeatMasker.

### Genome annotation

Each of the three genomes was independently annotated using Maker v2.31.10 [27], following the protocol given in [28]. In short, Maker is usually run several times successively and uses the gene models generated in one round to train *ab initio* gene-predictors and improve the initial gene models in the next round (see below). We used the above-mentioned hardmasked genomes and carried out their annotation using the proteomes of three closely-related species, namely *Pararge aegeria* [29], *Maniola hyperantus* [30] and *Bicyclus anynana* [31]. For each species, the output files were merged into a gff3 file that was then used to generate the necessary files to train SNAP (version 2006-07-28), an *ab initio* gene finding program [32]. A second run of Maker with the above-mentioned gff3 file and the .hmm file provided by SNAP resulted in a second gff3 file that was used to train SNAP a second time. A third round of Maker with the second gff3 and .hmm files was followed by the training of Augustus (3.3.3), another gene prediction tool [33], with the third gff3 file. A final round of Maker with the third gff3 file and the files generated by Augustus led to the fourth and last gff3 file, containing all the genome features for each species.

Protein-Protein BLAST 2.9.0+ (-evalue 1e-6 -max_hsps 1 -max_target_seqs 1) was then used to assess putative protein functions in each *Morpho* species by comparing the protein sequences given by Maker to the protein sequences from the annotated genomes of *Maniola jurtina* [29], *P. aegeria* [29], *B. anynana* [31] and *S. littoralis* specifically for the detection of OR sequences [34]. We used BUSCO to assess the completeness of the proteome with the protein mode and the *lepidoptera_odb10* database on the annotated gene set produced by MAKER [24].

### Phylogenetic analysis

To specifically compare the exon sequences of the opsins detected in the *Morpho* genomes to the opsins described in other Lepidoptera, we retrieved the coding sequences of opsins from NCBI and used the software Mega v.11 [35] to build a maximum likelihood tree and compute the associated bootstrap values.

Regarding the OR repertoire in the three *Morpho* species, we curated the sequences obtained by blast comparison of the Maker-annotated genes on the reference genome of *S. littoralis*, as a number of sequences showed incorrect lengths (< 300 or > 500 amino acids). We used exonerate version 2.4.0 [36] with the options -maxintron 2000 independently in each *Morpho* species. The exonerate alignment files and the assemblies were used with InsectOR (http://caps.ncbs.res.in/gws_ors/), a website specifically designed to help predict OR genes from insect genomes, with the option HMMSEARCH against 7tm_6 [37], [38]. The sequences uncovered with insectOR for each *Morpho* species were aligned with the ORs of *S. littoralis* using MAFFT [39] and we generated a maximum-likelihood phylogenetic tree with IQ-TREE version 2.2.0 [40] with the options -bb 1000 and -nt AUTO.

### Synteny and rearrangement detection

To assess variation in chromosome-scale synteny, we compared the assemblies of each *Morpho* to the assembly of *M. jurtina*, the closest relative of *Morpho* with a karyotype of 29 chromosomes and for which a high quality chromosome-level assembly (based on N50 values and Busco score, accession ID GCF_905333055.1) is available [29]. We used MUMmer 3.23 [41] to align the masked assembled genomes of *M. helenor, M. achilles* and *M. deidamia* to the *M. jurtina* genome. The output produced by MUMmer is an ASCII delta file that was then filtered and parsed using the utility programs delta-filter and show-coords from MUMmer. Synteny was visualized with the MUMmer results in R with the packages circlize v 0.4.12 [42] and Paletteer (https://github.com/EmilHvitfeldt/paletteer) using the Rscript from [43] described here: https://github.com/bioinfowheat/Polygonia_calbum_genomics/blob/7c75aac624157faa3ab229e3fc1e0e315302194d/synteny/circlePlot/nucmerOutput.R, removing short contigs, short alignments (less than 200bp) and low identity alignments (less than 90% identity).

In order to detect potential genome rearrangements between *Morpho* and closely-related species, we estimated the whole-genome collinearity between the *Morpho* assemblies and five closely-related Nymphalidae species whose genomes exhibit a good-quality assemblies in the NCBI genome database: *M. jurtina* (GCA_905333055.1), *P. aegeria* (GCA_905333055.1), *Erebia ligea* (GCA_923060345.2), *Melanargia galathea* (GCA_920104075.1) and *Lasiommata megera* (GCA_928268935.1) using D-GENIES [44]. Paired alignments between a *Morpho* species and one Nymphalidae species were performed using the minimap2 aligner [45] in D-GENIES, treating each *Morpho* species genome as the query and the Nymphalidae species genome as the target reference. We also used D-GENIES to pair-compare the genomes of the three *Morpho* species. As D-GENIES revealed differences between *Morpho* species in the contig corresponding to the Z chromosome (see results), we used SyRI [46] to study in detail the rearrangements in the sequences of this contig between the three species. We generated paired alignments of the Z contig with minimap2 and ran SyRI with the option -c on .sam files. SyRI requires that the two compared genomes represent the same strand and in the case of *M. achilles*, the orientation of the sequence produced by HiFiasm was the complementary to the sequences of *M. helenor* and *M. deidamia*. We then reverse-complemented this sequence in order to make the alignments. All the genomic structures predicted by SyRI were plotted using plotsr [47].

## Results

### Comparing karyotypes between species

First, we characterized the karyotypes of the three studied species (see Supplementary Fig. 2 to visualize the chromosomes). In *M. helenor*, the detected number of diploid chromosomes ranged from 54 to 56 in the different replicates of mitoses, with a discreet mode at 2n = 56. This variation is probably due to technical difficulties. The presence of n =28 bivalents in metaphase confirmed the diploid number of 2n = 56 chromosomes. In *M. achilles*, four specimens had the same modal chromosome counts: mitoses: 2n = 56 chromosomes; pachynema: n = 28 bivalents; Metaphases I: n = 28 bivalents; Metaphases II: *n* = 28 chromosomes with 2 chromatids. Surprisingly, the karyotype of the last male was quite different, with a modal number of 84 mitotic chromosomes. Interestingly, there was the same number (n = 28) of elements as above at the pachynema stage, indicating that they were trivalents. They were thicker than bivalents and a more careful analysis showed the recurrent asynapsis of one of the 3 chromosomes (Supplementary Fig. 3). No “normal” metaphase I or II was observed. It was concluded that this specimen was triploid with 3*n* = 84, and probably sterile. In *M. deidamia*, the diploid chromosome number had a discreet mode of 2*n* = 54, suggesting a slightly smaller number of chromosome pairs (n = 27) in this more distantly-related species. Our result are consistent with the modal number of chromosomes in the Morphinae (*n* = 28) described in previous karyotypic studies conducted in 8 Morpho species [48], where the reported number of chromosomes was also n = 28 for both *M. helenor* and *M. achilles*.

GenomeScope analyses suggested very high levels of heterozygosity for the three species (Table 1). In all of them, the N50 and contig sizes were generally larger in the assemblies produced by Hifiasm than in IPA and Flye assemblies (see supplementary table 1). The BUSCO scores revealed a very high percentage of duplicated sequences, especially in the assemblies produced by IPA and Flye. The use of purge_dups strongly reduced the number of duplicates, the estimated size of the genome and the number of final contigs (see Supplementary Fig. 4 and Sup. Table 1). Hifiasm and the post treatment with Purge_dups v1.2.5 gave an assembly of 143 contigs for *M. helenor* (size of the longest contig: 42411663 bp), of 32 contigs for *M. achilles* (size of the longest contig: 24854087 bp) and of 58 contigs for *M. deidamia* (size of the longest contig: 22518629 bp) (sup. Table 1). The Rebaler pipeline identified a circular mitochondrial genome of 15,336 bp for the species *M. helenor*, 15,340 bp for *M. achilles* and 15,196 bp for *M. deidamia*.

#### Annotation of repetitive region

In each of the three species of *Morpho*, we annotated around 50% of the genome as repeated elements (Supplementary Fig. 5). In *M. helenor*, 241,166,073 bp (51.28% of the genome) corresponded to repeated elements, 261,488,514 bp (54.65% of the genome) in *M. achilles* and 255,779,512bp (52.75% of the genome) in *M. deidamia*. The repetitive elements categories are shown in Supplementary Fig. 5. For the three species, long interspersed nuclear elements (LINE’s) accounted for the largest percentage (between 13.53% and 17.22%) of the repeated elements in the genomes.

### Genome annotation

We recovered 12,651, 12,978 and 12,093 protein-coding genes in the genomes of *M. helenor*, *M. achilles* and *M. deidamia* respectively. These values are comparable to what was found in *Maniola hyperantus* (13,005 protein-coding genes) and *P*.

**Table 1.** Genome heterozygosity estimated with GenomeScope and Genome statistics for the assemblies of three *Morpho* species using different computational methods. Assemblies were purged using purge_dups. Statistics were obtained with BBMap. The assembly produced with Hifiasm for the individual M. *deidamia* was not purged with purge_dups as BUSCO results on the preliminary assembly revealed a very low duplicate content.

|  | *M. helenor* | *M. achilles* | *M. deidamia* |
| --- | --- | --- | --- |
| Heterozygosity (%) | 3.35 | 2.78 | 1.68 |
| Assembly method |  |  |  |
| Hifiasm |  |  |  |
| Total contigs | 143 | 32 | 58 |
| Genome size | 470.254 Mb | 478.514 Mb | 489.914 Mb |
| N50 | 12 Mb | 12 Mb | 13 Mb |
| IPA |  |  |  |
| Total contigs | 128 | 56 | 47 |
| Genome size | 473.620 Mb | 493.177 Mb | 481.177 Mb |
| N50 | 17 Mb | 14 Mb | 13 Mb |
| Flye |  |  |  |
| Total contigs | 134 | 114 | 291 |
| Genome size | 466.515 Mb | 477.638 Mb | 484.463 Mb |
| N50 | 21 Mb | 20 Mb | 34 Mb |

*aegeria* (13,515 protein-coding genes), but were lower than in *M. jurtina* (13,777 protein-coding genes) and *B. anynana* (14,413 protein-coding genes). Busco results for the proteome and transcriptome are presented as supplementary material (see Supplementary Fig. 5 and 6) In order to assess if the annotations were complete, we estimated in each species the percentage of proteins with a Pfam domain as this value has been found to vary between 57% and 75% in eukaryotes [49]. This value ranged from 65,50% in *M. achilles* to 71,32% in *M. helenor* with an intermediate value of 70,42% in *M. deidamia*, thus showing that the annotations were of good quality. Proteome completeness using BUSCO was also high. From a set of 5286 single-copy orthologues from the lepidoptera lineage, the proteome completeness varied between 69% and 79% depending on the species (Supplementary Fig. 5). We were thus able to further investigate gene families that could be involved in pre-zygotic isolation through duplication or loss events. This includes genes having a role in vision (*L-opsin*) but also chemosensory genes such as odorant and gustatory receptors that reflect the degree of species specialization.

#### Duplications in opsins genes

Vision in butterflies notably relies on opsins, for which three major types of molecules have been described depending on their wavelength of peak absorbance: in the ultraviolet (UV, 300-400 nm), blue (B, 400-500 nm) and long wavelength (L, 500-600 nm) part of the visible spectrum. Opsins are encoded by UV, B and L opsin genes. We investigated the number of copies for each opsin gene in the three *Morpho* species. We consistently found one copy of the UV opsin gene and two copies of the B opsin genes in the three *Morpho* species. Duplications of the L-opsin were observed in *M. achilles, M. deidamia* and *M. helenor*. In the other reference genomes *M. jurtina, B. anynana* and *P. aegeria*, a single copy of the UV opsin gene, the B opsin gene and the L opsin gene were found. By comparing the L-opsin sequences using a maximum likelihood tree based on the exon sequences (Fig. 2), we showed that the duplications observed in *Morpho* butterflies probably occurred independently from previously described duplications that happened in other clades of Lepidoptera. The phylogenetic relationships between the copies in the three species reveal that the duplications observed in the three *Morpho* species probably occurred before their speciation (Fig. 2). The detection of the different copies in different species within the Morpho genus and in closely-related genus is now required to precisely characterize the evolutionary origin of these duplications.

**Figure 2.**
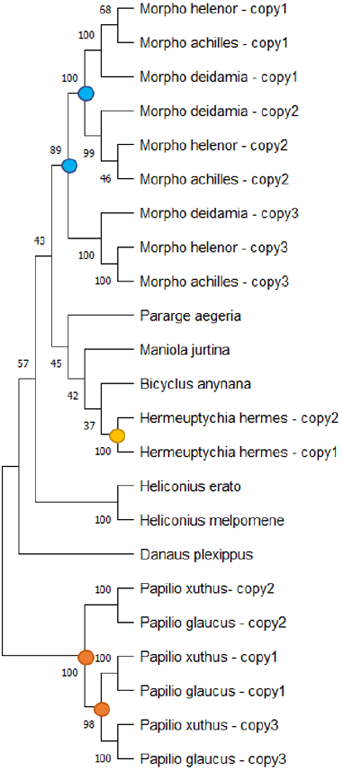
Maximum Likelihood tree of L-opsin exon sequences detected in the genomes of *M. helenor, M. achilles* and *M. deidamia* and other butterflies species, with bootstrap values. The colored dots indicate the putative locations of the duplication events on the tree: the putative origin of duplications of the L-opsin observed within the genus *Morpho* appear in blue, while the duplications that occured in the *Hermeuptychia hermes* clade and in the *Papilio* clade appear in yellow and orange respectively

#### Odorant and gustatory receptors

In order to estimate the number of *OR* and *GR* genes in the three *Morpho* species, we blasted our Maker-annotated genes on the reference genome of *Spodoptera littoralis*. In this moth species, 60 *OR* and 16 *GR* genes were curated [50]. Interestingly, we recovered only 31 *OR* genes including Orco in *M. helenor*, 32 in *M. achilles* and 36 in *M. deidamia*, while we found 14 *GR* genes in *M. helenor* and 16 in *M. achilles* and *M. deidamia*. With insectOR, we found 36 OR genes including Orco in *M. helenor*, 37 in *M. achilles* and 38 in *M. deidamia*, confirming the major loss of ORs in our three Morpho species. For comparison we blasted against the same reference genome of *S. littoralis* the annotated sequences of the three outer Lepidopteran species used in the previous analyses and uncovered a much higher number of *OR* and *GR* genes with 61 *OR* and 28 *GR* in *M. jurtina*, 60 *OR* and 35 *GR* in *B. anynana* and 50 *OR* and 20 *GR* in *P. aegeria* respectively. The drastic reduction of chemosensory receptors, particularly in the number of *OR* genes in the three *Morpho* species could potentially reflect a higher degree of specialization to their respective biochemical environment. A phylogenetic analysis of *Morpho* ORs along with those of *S. littoralis*, the sole Lepidopteran species for which a considerable number of ORs were functionally deorphanized and divided into three chemical classes (aromatics, terpenes and aliphatics) as described in [34], showed that the loss of ORs in *Morpho* were not clustered around a particular set of genes (Supplementary Fig. 9). Further functional characterization coupled with precise ecological investigations are therefore needed to understand the loss of ORs in the *Morpho* genus.

### Synteny and rearrangement detection

#### Conserved synteny with other Lepidoptera species

We found a high concordance between the *n* = 29 chromosomes of *M. jurtina* and the contigs of the three *Morpho* species (Fig. 3). The MUMmer alignment and the post alignment treatment to remove short contigs and low identity alignments reduced the assembly to 27 contigs containing 97% of the total genome for *M. helenor* (removing 117 short contigs from the original assembly), 29 contigs (98% of the genome) for *M. achilles* (3 contigs removed) and 27 for *M. deidamia* (31 contigs removed) (Fig. 3).

**Figure 3.**
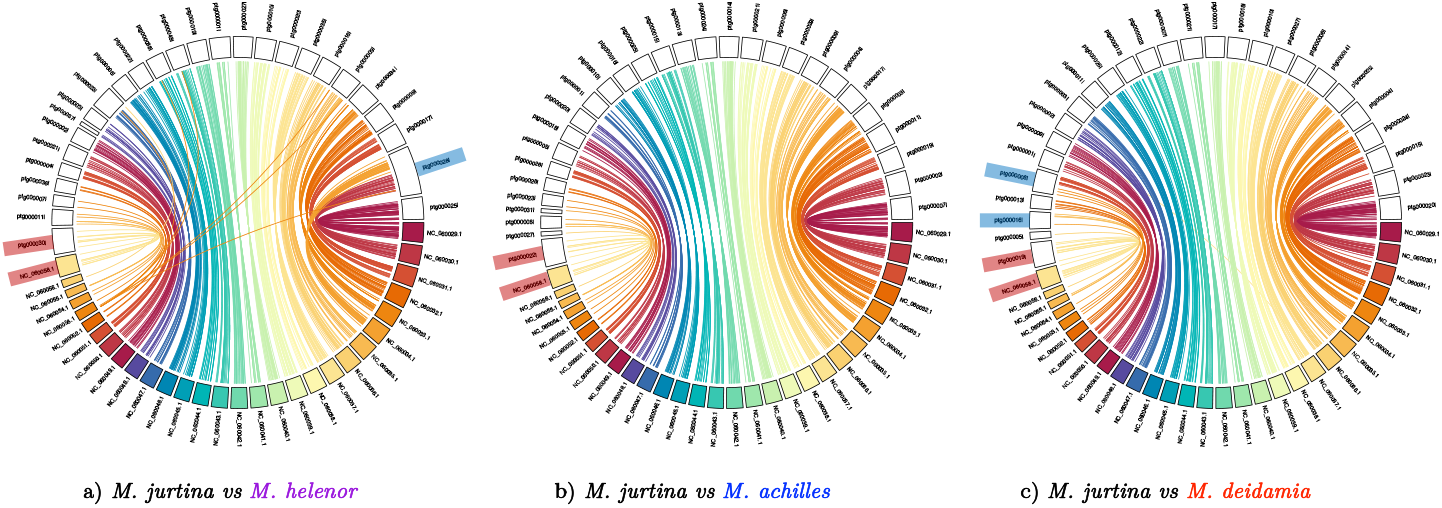
Synteny between the chromosome-assembled genome of *Maniola jurtina* (colored chromosomes) and the genome assemblies of the species *Morpho helenor* (a),M. *achilles* (b) and *M*. *deidamia* (c). Equivalent chromosomes/contigs are linked by same color ribbons. Chromosome Z for each species is labeled in red. Single chromosomes in *Morpho* that are not assigned to a single chromosome in *M. jurtina* are labeled in blue

The synteny plot between *M. helenor* and *M. jurtina* showed 27 contigs for *M. helenor*, one contig less than expected based on its karyotype of n=28. In the plot, one single contig (ptg000028l) was assigned to two different chromosomes from the *M. jurtina* assembly (chromosomes 2 and 6, NC_060030.1 and NC_060034.1). Contig ptg000028l is twice the size of any other contig found in the three *Morpho*
species analyzed here. Based in the differences between the number of contigs and the karyotype and the unexpectedly big size of the contig ptg000028l, we believe the difference in chromosome number between *M. jurtina* and *M. helenor* can be explained by an overassembly of the genome of *M. helenor* by hifiasm, that assigned one single contig to two different chromosomes from the *M. jurtina* assembly (Fig. 3). In *M. deidamia*, the Hifiasm assembly showed a single contig ptg000008l (size 20.29 Mb) containing chromosomes NC_060051.1 and NC_060052.1 (sizes 10.05 Mb and 9.43 Mb respectively) from *M. jurtina*. Because the number of contigs recovered for this species is in accordance to the karyotype of n=27, the differences between *M. deidamia* and *M. jurtina* suggest that in this case chromosomes NC_060051.1 and NC_060052.1 in *M. jurtina* may have fused to form contig ptg000008l in *M. deidamia*. Other rearrangement in this species compared to *M. jurtina* seems to be the contig ptg0000161l, that appears to contain small portions of chromosomes NC_060054.1 and NC_060055.1 from *M. jurtina*.

For the three *Morpho* species, we were able to identify a single contig corresponding to the chromosome Z (NC_060058.1) in *M. jurtina* (contig ptg000030l in *M. helenor*, contig ptg000024l in *M. achilles* and contig ptg000019l in *M. deidamia*).

We also found a high level of colinearity between the genomes of the three *Morpho* species and the five Nymphalidae species used for comparisons. The alignment between *M. jurtina* and the three *Morpho* species (Fig. 3) was very similar to the alignments obtained for the other Nymphalidae (Supplementary Fig. 6) and confirmed that the assembly of the genome of *M. helenor* by hifiasm might have merged together two chromsomes: the single contig ptg000028l was scattered into two chromosomes in the other Nymphalidae. Although collinearity was generally high, we detected some putative inversions located in regions that varied among pairs for the three *Morpho* species in comparison with the Nymphalidae (see Supplementary Fig. 6). Interestingly, the contig corresponding to the chromosome Z was the only one consistently showing inversions in the pairwise genome-wide alignments (see Supplementary Fig. 6).

#### Inversions in the Z-chromosome between the three sympatric *Morpho* species

Pairwise whole genome alignments of the three *Morpho* species showed a very high similarity between genomes (see Supplementary Fig. 7). The only contig that differed between species was the one corresponding to the Z chromosome. SyRI identified one inversion of 1.6 Mb between *M. helenor* and *M. deidamia*, five inversions (comprising one of more than 1.8 Mb) between *M. helenor* and *M. achilles* and two between *M. deidamia* and *M. achilles* with one of 1.6 Mb (Fig. 4). Interestingly, the inversion found in *M. deidamia* when compared to *M. achilles* or *M. helenor* has the same size and is located in exactly the same position of the chromosome (from bp 1567583 to 3192401), suggesting that this inversion is ancestral to the speciation of *M. achilles* and *M. helenor*. In the case of *M. achilles vs. M. helenor* two inversions were found flanking the site of the putative ancient inversion and a bigger inversion was found at the end of the chromosome (Fig. 4).

**Figure 4.**
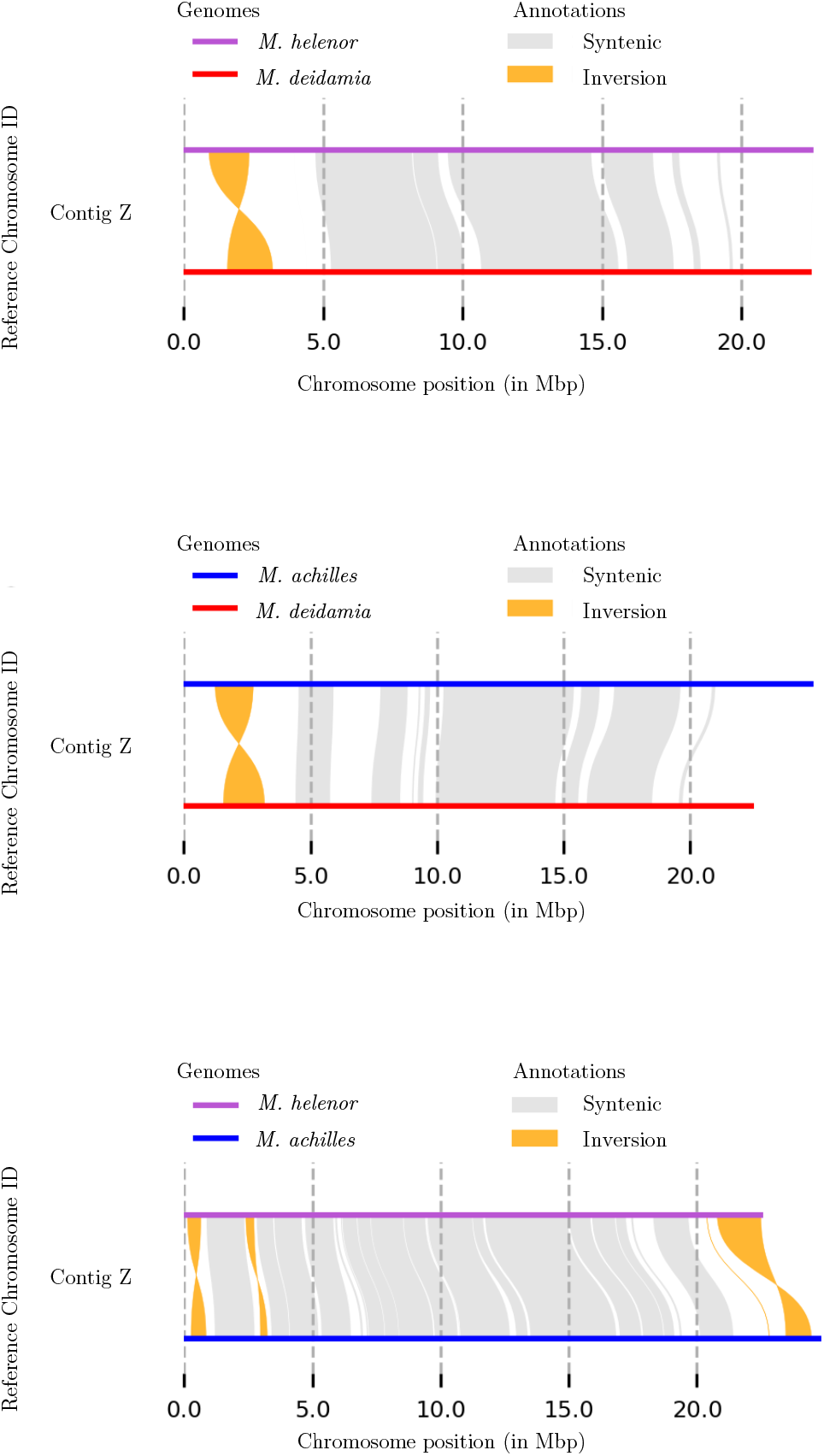
Synteny and rearrangement (SyRI) plot of the paired comparisons for the Z contig between the three *Morpho* species. Upper figure: *M. helenor* and *M*. *deidamia;* middle: *M*. *deidamia* and *M*. *achilles;* lower: *M. helenor* and *M*. *achilles*.

## 1 Discussion

### Assembly of heterozygous Lepidoptera genomes with a high proportion of repeated elements

We generated *de novo*, reference-quality genome assemblies for three emblematic species of Amazonian butterflies: *M. helenor, M. achilles* and *M. deidamia*. Our results indicate genome sizes comprised between 470 Mb and 489 Mb, similarly to most of the closely-related Nymphalidae species sequenced so far, *e.g. B. anynana* (475 Mb), *P. aegeria* (479 Mb) or *M. jurtina* (429 Mb). This is also close to the 479 Mb estimated from phylogenetic comparison using the taxon-centred database “Genomes on a Tree” (GoaT) [51]. The final number of contig within each of the three species ranged from 27 to 29, close to the number of chromosome pairs observed in our cytogenetics study. The numbers of chromosomes found in those French Guiana samples (i.e. in the subspecies *M. helenor helenor* and *M. achilles achilles*) is consistent with those found in other subspecies of both species in previous studies [48]. The available sequenced species of Nymphalidae that are closely-related to the genus *Morpho* also generally show 29 pairs of chromosomes (28 autosomes, plus Z and W sex chromosomes), which is close to the chromosomal numbers observed in the three *Morpho* species studied here. The mapping between the assemblies of *Morpho* species to the chromosome-level assembly of *Maniola jurtina* and the post-treatment to eliminate small contigs allowed us to identify between 27 and 29 contigs in *Morpho* that were homologous to *Maniola jurtina* chromosomes, including the contig corresponding to the *Z* chromosome. This suggests a high conservation of chromosomal synteny among closely-related Nymphalidae species, which is consistent with the high level of synteny observed throughout the whole Lepidoptera clade [52]. In the three species, genome heterozygosity was very high (from 1.68% in *M. deidamia* to 3.35% in *Morpho helenor*) and heterozygosity presents a major challenge in *de novo* assembly of diploid genomes. Indeed, levels of heterozygosity of 1% or above are considered “moderate to high” and most assemblers struggle when two divergent haplotypes are sequenced together, as heterozygosity may impair the distinction of different alleles at the same locus from paralogs at different loci [53]. Then, final assemblies of heterozygous genomes are expected to be of poor-quality, highly fragmented and containing redundant contigs [54]. Hifiasm generated the most completely haplotype-resolved assemblies, nevertheless the level of heterozygosity clearly impacted the quality of the assemblies and a post treatment to remove duplicated sequences was necessary for the two most heterozygous genomes (*M. helenor* and *M. achilles*), showing the difficulty that heterozygosity still imposes to long-read heterozygosity-aware assemblers. Such a high heterozygosity has been observed in other genomes of Lepidoptera [31] and can be a signature of high effective population sizes. The wide Amazonian distribution of these species, and their flight activity could contribute to such high level of genetic diversity within population, because elevated dispersal contribute to increase gene flow within each species throughout their geographic range. Our results also showed that around 50% of the genomes of the sequenced *Morpho* was composed of repeated elements, a very high proportion as compared to other genomes of Lepidoptera. In Lepidoptera, TE content has been found to be correlated with genome size [55], but in the case of the three *Morpho* species studied here, the repeat content is higher than for other species with similar genome sizes such as the *Bombyx mori* moth, with a genome size estimated at 530 Mb and a TE content of 35% [56] or the more closely-related species *Bicyclus anynana* with a genome size of 475 Mb and a repeat content of 26% [31].

### Structural variations between genomes of sympatric species

The karyotype and assembly analyses suggest some differences in chromosome number between the three sympatric *Morpho* species studied here, particularly between *M. deidamia* (27 chromosome pairs) *vs M. helenor* and *M. achilles* (28 chromosome pairs). Differences in chromosome numbers and other chromosomal rearrangements may strongly affect reproductive barriers. Two groups of models have been proposed to explain how chromosomal rearrangements prevent gene-flow and contribute to species maintenance and speciation. First, hybrid-sterility models suggest reduced fertility or viability in individuals heterozygous for chromosomal rearrangements. These models are considered to be inconsistent and difficult to evaluate [4]. More recently, suppressed-recombination models propose that chromosomal rearrangements permit speciation in sympatry because they reduce recombination between chromosomes carrying different rearrangements [4]. Indeed, in Lepidoptera, differences in chromosome number are proposed to be an important mechanism leading to species diversification in *Agrodiaetus, Erebia* and *Lysandra* butterflies ([57–59]).

Besides differences in chromosome numbers, we systematically found inversions in the contig corresponding to the Z chromosome when comparing the genomes of *Morpho* to the other Nymphalidae and between the three different *Morpho* species. Inversions are also a type of chromosomal rearrangement known to occur throughout evolution and are considered an important mechanism for speciation particularly for species living in sympatry ([1, 4]). Empirically and theoretically, it has been suggested that inversions may have contributed to speciation in sympatry in different groups of animals. In two ascidians species of the genus *Ciona* and in insects like *Drosophila* inversions may promote speciation by reduction of the fitness or by causing sterility of heterozygotes ([60, 61]). In the *Anopheles gambiae* species complex, inversions may allow for ecotypic differentiation and niche partitioning leading to different sympatric and genetically isolated populations [62]. In groups like paserine birds where sexual differentiation is controlled by a ZW sex chromosome system (females being the heterogametic sex), inversions in the Z chromosome in particular seem to explain speciation in sympatry between close species. Cytological data show that across the Passeriformes, the Z chromosome has accumulated more inversions than any other autosome and that the inversion fixation rate on the Z chromosome is 1.4 times greater than the average autosome. Interestingly, inversions on the Z chromosome are significantly more common in sympatric than in allopatric closely related clades ([63, 64]).

In Lepidoptera, the role of inversions in speciation in sympatry has been studied in the species *Heliconius melpomene* and *H. cydno*, two sympatric species that can hybridize (although rarely) in the wild [65]. The analyses of the genomic differences between the two species showed some small inversions (less than 50 kb) and there was no evidence for a reduction of recombination in hybrids, suggesting that in this case, inversions were not involved in the maintenance of the species barriers and other processes such as strong mate preference could prevent hybridisation in the wild [65]. In the *Morpho* species studied here however, we found inversions between *Morpho* Z chromosomes that were longer than 1.5 Mb. Models suggest that to be associated with adaptive traits or species barriers, inversions should typically be megabases long in order to be fixed in populations [65]. The position of the inversion in the Z contig when comparing *M. helenor* or *M. achilles* to *M. deidamia* is at the exact same place in *M. deidamia*’s genome, suggesting that this specific inversion likely occurred before the speciation between *M. achilles* and *M. helenor*. When comparing *M. helenor* to *M. achilles*, we found two different smaller inversions that are not found in *M. deidamia* and that are close to the putative ancestral inversion region, suggesting that these two smaller inversions could have appeared after the speciation between *M. achilles* and *M. helenor*. At the moment, we do not know if the inversions segregate at different frequencies in the *Morpho* populations or if they are fixed. Population analyses are needed to answer this question and to enlighten what evolutionary forces could be acting to maintain them. The copy number variation detected in genes involved in colour perception (i.e. *L-opsin*) may also play a significant role in reproductive isolation in these sympatric species. For instance, the three copies of L opsins found in the Papilio genus (Fig. 2) have been found to also show subfunctionalization and neofunctionalization [66]. The duplication followed by genetic divergence observed in these three mimetic *Morpho* species may improve their visual discrimination capacities, and facilitate species recognition, therefore reinforcing barriers to gene flow in sympatry. Genes potentially involved in colour pattern variations (e.g. *bric – a – brac* or *bab*) may also play a role in prezygotic isolation but they were not thoroughly investigated here as their functional evolution involves changes in regulatory sequences rather than events of duplication or gene loss [67]. Interestingly, an orthologous search of the putative proteic sequences of each *Morpho* species against those of *M. jurtina* allowed us to uncover different copy numbers of the gene *bric – a – brac*, which play a significant role in differences of UV iridescence between males of two incipient species of sulphur butterflies [68]. The copy responsible for the presence/absence of UV iridescence is located on the Z chromosome and in the three *Morpho* species, we found one or more copies of *bric – a – brac* on the contigs that correspond to the Z chromosome: *M. deidamia* had one copy of *bric – a – brac*, while *M. helenor* and *M. achilles* displayed two copies of this gene. It seems however that the second copy in *M. helenor* and *M. achilles* correspond to truncated copies of *bric – a – brac*. While this is certainly the sign of an ancient duplication followed by a pseudogenization event, this could lead to further investigations of putative functions of the truncated copies. It is worth noting that variations in the number of *bab* copies was also observed in the three reference genomes used for the blast: *M. jurtina* had two copies on the Z chromosome (including a truncated copy), *B. anynana* had only one and *P. aegeria* had none. The investigation of gene levels of polymorphism on the Z chromosomes would also be of great interest as genes linked to the Z chromosome are often among the most divergent between closely-related species [69].

Altogether, the assembly and annotation of these three mimetic species of *Morpho* butterflies reveal differences in chromosome numbers, the presence of several Mb-long inversions in the Z chromosome, as well as copy number variation and genetic divergence among copies of genes that may play a significant role in reproductive isolation. Our study thus open new avenues into the investigation of the ecological and genomic factors involved in sympatric speciation and its reinforcement.

## Acknowledgements

The authors would like to thank Patrick Blandin for his continuous support on our *Morpho* studies. They also thank Mélanie McClure, Mathieu Chouteau, Camille Le Roy and Ombeline Sculfort for their help during field work in French Guiana. We thank Elise Gay, Romuald Laso-Jadart, Pierre Lesturgie, Christelle Fraisse, Clément Gilbert and Quentin Rougemont for help with some scripts and for discussions of previous versions of the manuscript. We thank Charlotte J. Wright and Niklas Wahlberg for their careful reading of our manuscript and their many insightful comments and suggestions. We are thankful to the Plateforme de Calcul Intensif et Algorithmique PCIA, Muséum national d’histoire naturelle, Centre national de la recherche scientifique, UAR 2700 2AD, CP 26, 57 rue Cuvier, F-75231 Paris Cedex 05, France and the Genotoul bioinformatics platform Toulouse Occitanie (Bioinfo Genotoul, https://doi.org/10.15454/1.5572369328961167E12).

