## Supplementary for "Genome assembly of three Amazonian *Morpho* butterfly species reveals Z-chromosome rearrangements between closely-related species living in sympatry"

### Supplementary material

March 1, 2023

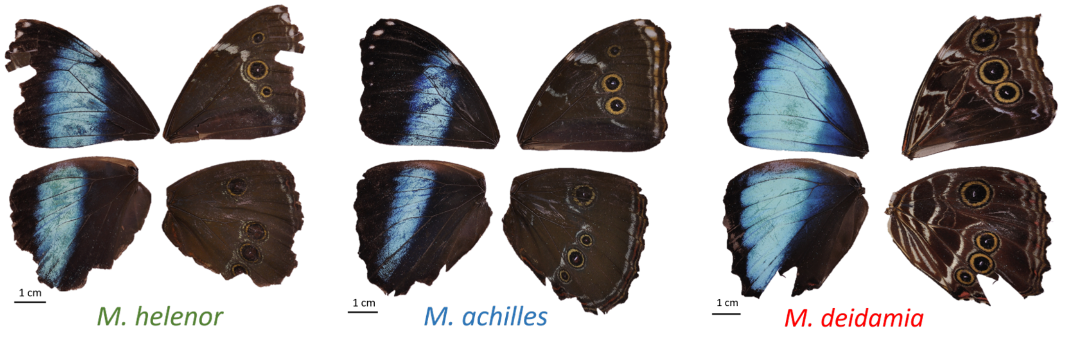


Figure 1: Pictures of dorsal and ventral sides of the wings of the sequenced specimens of *M. helenor*, *M.achilles* and *M.deidamia* sampled in French Guiana

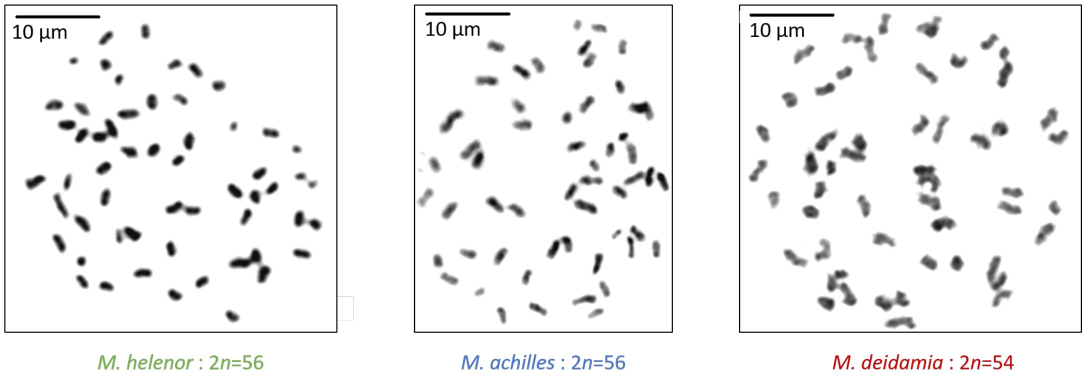


Figure 2: Pictures of caryotypes obtained from specimens of *M. helenor*, *M.achilles* and *M.deidamia* sampled in French Guiana, note that the chromosome number differ between *M. deidamia* and the other two species.



Draft

Table 1: Genome statistics for the assemblies of three *Morpho* species before the use of Purge_dups and using different assemblers. Statistics were obtained with BBMap.

| | *M. helenor* | *M. achilles* | *M. deidamia* |
|---|---|---|---|
| Before purge_dups | | | |
| Hifiasm | | | |
| Total scaffolds | 207 | 56 | 58 |
| Genome size | 547.058 Mb | 490.219 Mb | 489.914 Mb |
| N50 | 14 Mb | 13 Mb | 13 Mb |
| IPA | | | |
| Total scaffolds | 241 | 142 | 121 |
| Genome size | 922.833 Mb | 954.108 Mb | 900.010 Mb |
| N50 | 43 Mb | 30 Mb | 25 Mb |
| Flye | | | |
| Total scaffolds | 334 | 305 | 656 |
| Genome size | 921.488 Mb | 954.510 Mb | 959.961 Mb |
| N50 | 51 Mb | 44 Mb | 70 Mb |

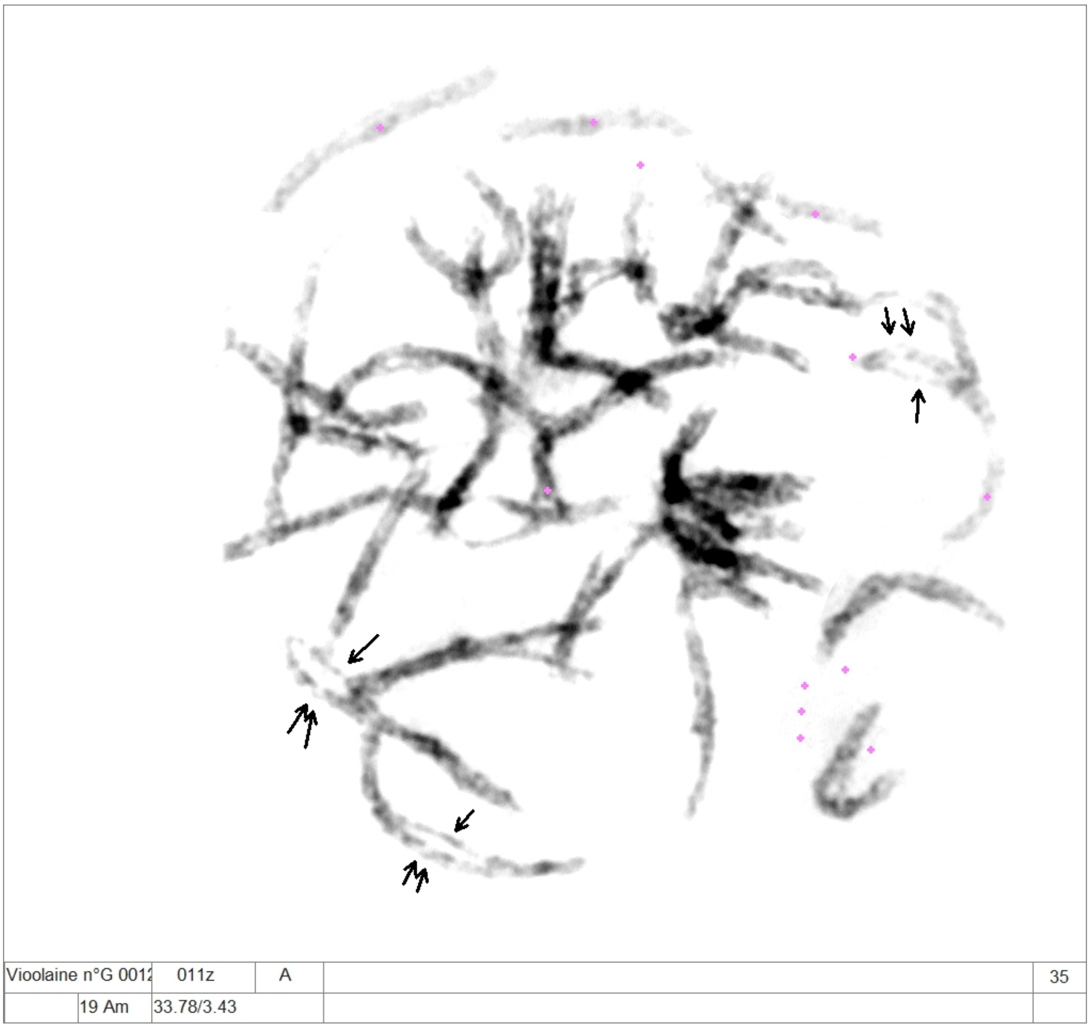


Figure 3: Picture of the putative triploid caryotype obtained from a specimens of*M.achilles*. There was the same number ($n = 28$) of elements as in the other *M. achilles* individuals at the pachynema stage. However, these elements were thicker than bivalents and a more careful analysis showed the recurrent asynapsis of one of the 3 chromosomes (see arrows).

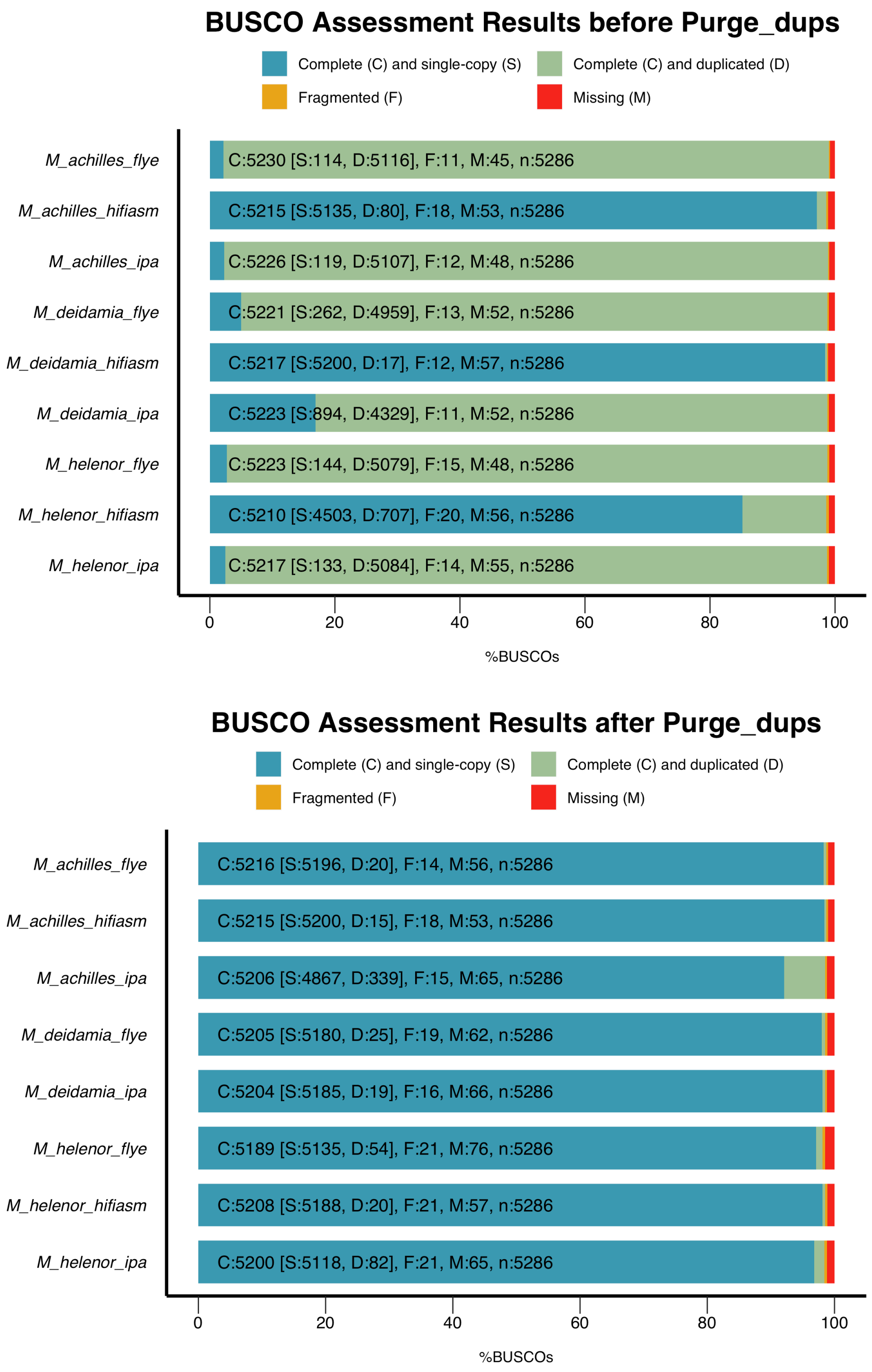


Figure 4: BUSCO assessment of the genome assembly completeness of *Morpho* species assembled using IPA, Flye and Hifiasm. Results correspond to the BUSCO score before (top) and after (bottom) the use of purge_dups

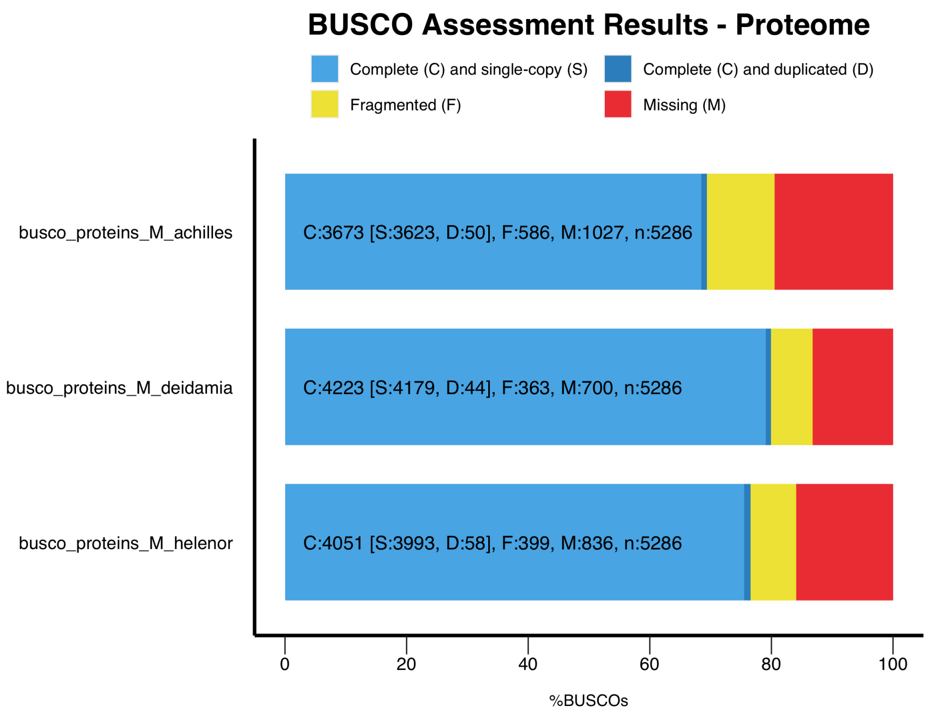


Figure 5: BUSCO analysis of gene annotation of *Morpho* species. BUSCO was run in "protein" mode on the annotated gene set produced by MAKER

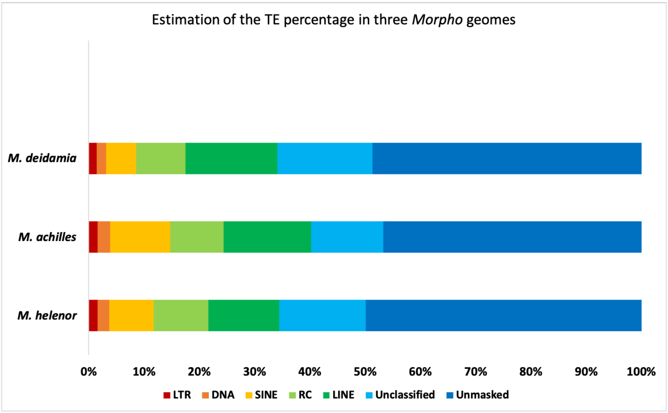


Figure 6: Proportion of each class of transposable elements (TEs) in the assembled genomes of the three *Morpho* species

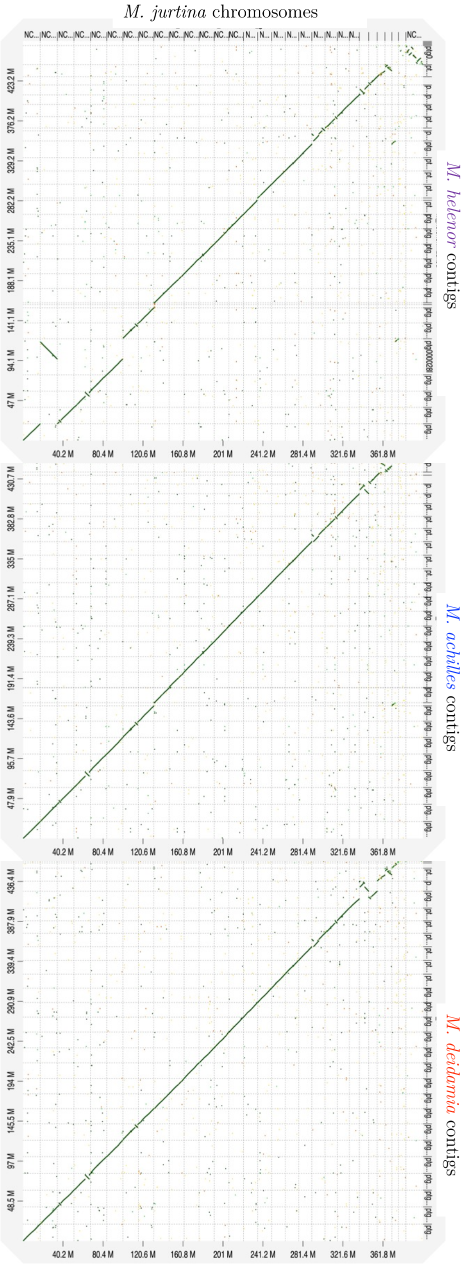


Figure 7: D-GENIES dot-plots comparing the genome assemblies of three *Morpho* species against the assembly of the species *Maniola jurtina*. Chromosomes of *M. jurtina* are in the X-axis while the Y-axis shows the scaffolds of (upper) *M. helenor* (middle),*M. achilles* and (bottom) *M. deidamia*. The alignment between the Z chromosome of *M. jurtina* and *Morpho* corresponds to the square in the upper right corner. Here we show only the plot between *Morpho* and *M. jurtina*, comparison with four other species of Nymphalidae are available upon request

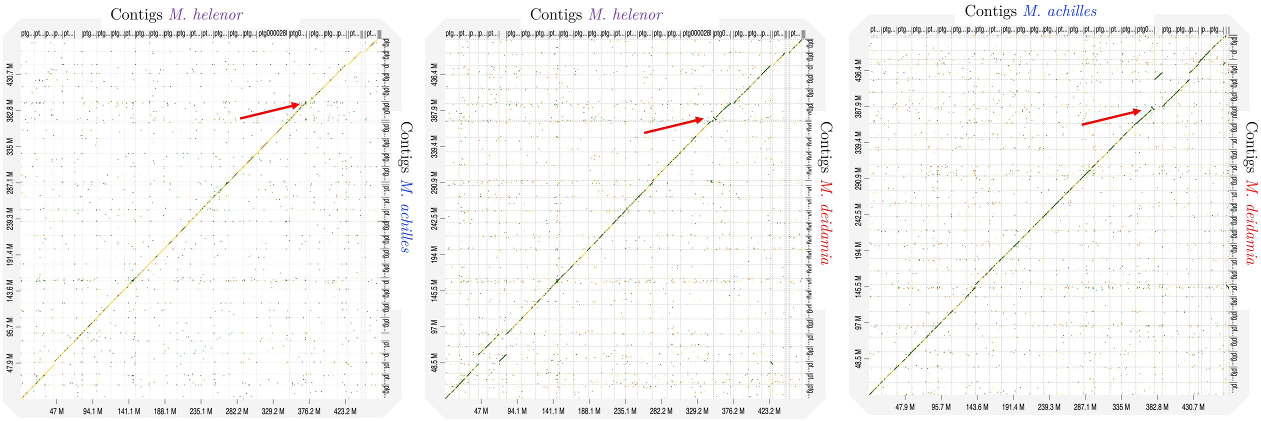


Figure 8: D-GENIES dot-plots comparing the genome assemblies of pairs of *Morpho* genomes. The red arrows indicate the inversions (in the scaffolds corresponding to the Z chromosome)

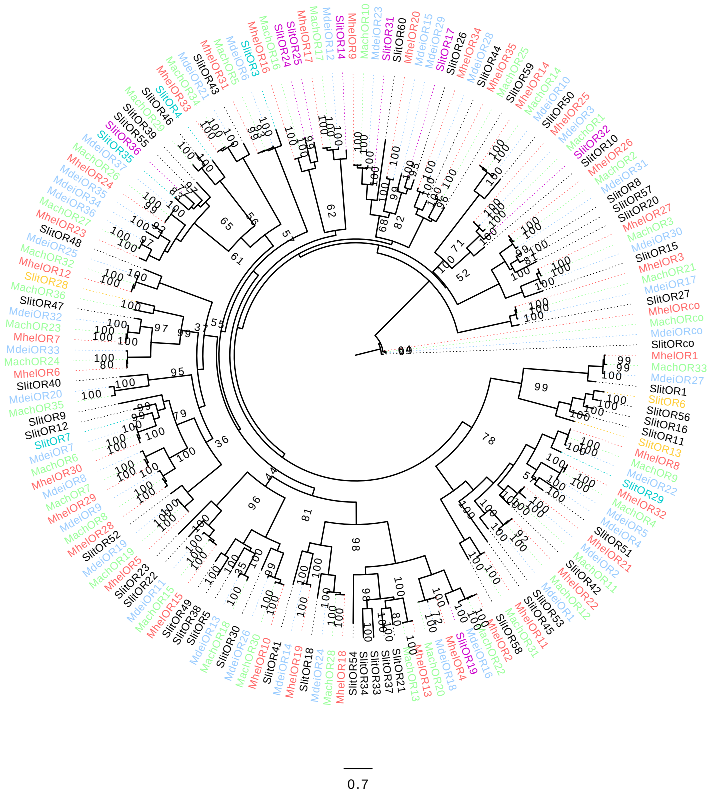


Figure 9: Phylogenetic tree by maximum likelihood of ORs built from amino-acid sequences of *M. helenor* "MhelOR" (red), *M. achilles* "MachOR" (green), *M. deidamia* "MdeiOR" (blue) and *S. littoralis* "SlitOR". The latest were colored following their chemical class (magenta: aromatics; cyan: terpenes; orange: aliphatics; black: unclassified) as defined in [**?** ].
